# Inferring EMT dynamics from cell cycle profiles using a hidden Markov framework

**DOI:** 10.1101/2025.10.28.685248

**Authors:** Annice Najafi

## Abstract

The Epithelial-to-Mesenchymal Transition (EMT) which is a driver of metastasis and a contributor to wound healing, embryogenesis, and trophoblast differentiation is highly context specific, making it difficult to infer robustly from sequencing data. Existing methods often require that epithelial, hybrid, and mesenchymal states all be present in the same dataset and rely on simplifying assumptions that limit generalizability. By contrast, cell cycle stage is routinely and reliably inferred from transcriptomic profiles. We leverage this robustness to recover EMT dynamics when direct EMT inference fails. Specifically, we learn an emission model that links latent EMT states to observed cell cycle stage frequencies using datasets with known EMT trajectories. We then use this model, together with the initial EMT distribution of a cell line, to demonstrate that we can reconstruct EMT trajectories from time-series measurements of cell cycle stage alone. Finally, we provide an open-source R package and a companion web application to make the approach accessible and reproducible.

**SIGNIFICANCE:** Elucidating EMT dynamics is central to understanding metastasis and therapy response, yet EMT is notoriously context-specific and often cannot be inferred reliably from transcriptomic data alone. We introduce a population-level hidden Markov model that links latent EMT states to observed cell cycle compositions via an estimated emission matrix thereby enabling the reconstruction of EMT trajectories and transition rates using only cell cycle fractions and a baseline EMT distribution (specific to the cell line). This leverages the strong coupling we observe between EMT progression and cell cycle regulation across cell lines, while remaining robust when direct EMT inference fails. We provide an open-source R package and web app to make these analyses accessible.

## 1 INTRODUCTION

EMT is a reversible and dynamic cell-state program that allows epithelial cells to acquire migratory and mesenchymal-like traits such as motility, invasiveness, and resistance to stress (1). Once viewed as a binary process, EMT is now increasingly recognized as a process with a number or continuum of intermediate phenotypes that can be broadly represented by three states: epithelial, hybrid, and mesenchymal (2). Transitions among these states underlie several fundamental biological processes such as embryonic development (3), wound healing (4), tumor progression (5), and trophoblast differentiation (6). At the same time, cell cycle progression which co-occurs with EMT orchestrates alternating phases of growth and division (7). The relationship between EMT and cell cycle progression is an important topic to study due to its implications for tumor dormancy, metastatic dissemination, and therapeutic resistance (8).

While some studies have shown that mesenchymal differentiation suppresses proliferation, leading to an enrichment of G1-arrested or quiescent cells (9, 10), this relationship has rarely been quantified systematically, either across time or across different cell lines. The recent advent of single-cell RNA sequencing allows us to study the coupling of these two processes in greater detail.

Observations suggest that the coupling between EMT and cell cycle regulation may be far more heterogeneous and context-dependent than previously appreciated (11). However, in many practical settings, EMT trajectories cannot be recovered directly, since existing EMT trajectory-inference methods often fail under noncanonical inducers, strong cell-line heterogeneity, or in the presence of stroma (12). Consequently, we need a model that infers EMT dynamics even when EMT signals are unreliable or missing by explicitly coupling a latent EMT process to the observed cell cycle composition and using the latter as an informative readout to reconstruct EMT transition rates and state trajectories from population-level data.

Previously, we developed a data-driven framework, COMET, to infer EMT trajectories from time-course single-cell RNA sequencing data (12, 13). We demonstrated that this approach can reliably estimate cell-state fractions and interstate transition rates from gene expression profiles of cell lines undergoing TGF*β*-induced EMT in vitro (12). However, the method struggled to accurately reconstruct EMT trajectories in cases where EMT was induced through other EMT induction factors such as EGF or TNF*α*. Interestingly, when we derived cell cycle fractions using the Seurat package in R (14), we observed a striking similarity in cell cycle dynamics across cell lines, even when EMT was triggered by distinct induction factors (12). This pattern suggested a deeper and possibly cell line-specific coupling between EMT progression and cell cycle regulation.

Here, we introduce a population-level Hidden Markov Model (HMM) that formalizes this coupling by linking unobserved EMT transitions to observed cell cycle distributions. In this model, EMT progression is described by a Continuous Time Markov Chain (CTMC), while the cell cycle phases constitute the emission layer. The emission matrix quantitatively characterizes how each EMT state expresses distinct cell cycle activity. When both EMT fractions and cell cycle distributions are available, the emission layer can be estimated using least squares or maximum-likelihood inference and can be used to reconstruct EMT trajectories in similar cases of a cell line undergoing EMT. In the results to follow, we first infer the emission matrix from data where both EMT and cell cycle fractions are available then use the HMM model to infer EMT trajectories from datasets where the EMT data is inaccessible. The method allows us to quantify the changes in interstate transition rates as a result of different EMT induction factor treatments. We conclude our study by providing an open-source R package (`hiddenCOMET`) for this model. This framework enables a unified, quantitative analysis of how EMT dynamics influence cell cycle stages across cell lines and treatment conditions and allows users to infer EMT dynamics in cases where EMT inference algorithms fail to reliably extract EMT state dynamics. Although, we introduce this method in the EMT and cell cycle context, the same methodology can be applied in similar scenarios where direct inference of a certain process is challenging due to context specificity.

## MATERIALS AND METHODS

### 1.1 Model Development

We use a CTMC model to track the phenotypic evolution of cancer cells as they undergo EMT. Our CTMC model is characterized by the following generator matrix similar to the one defined in Equation 3 of Najafi et al (12):

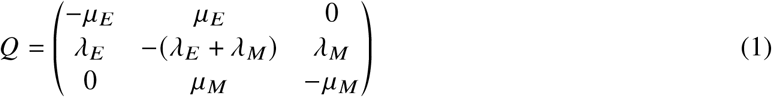

Here, the EMT process evolves via four transition rates that remain constant throughout the process (we note that unlike the original COMET framework, we do not separate the EMT process into different regimes or phases, and the process evolves via a single CTMC model. In our previous framework we also assumed symmetric transition rates into the hybrid state which is relaxed here. In addition, we note that some studies report phenotypic stability factors may not result in symmetric transition rates into the hybrid state (15) which further substantiates the use of a single regime model for simulating the EMT process. Consequently, we believe removing that assumption would expand the applicability of our framework). We take the matrix exponential of this generator matrix to find the transition probability matrix which evolves through time (*P* (*t* ) ) as follows:

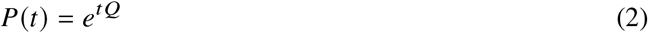

The transition probability matrix (3 × 3, corresponding to the three EMT states of epithelial, hybrid, and mesenchymal) determines the probability of transitioning from one state to another (the *P*_*i, j*_ (*t* )element of the transition probability matrix refers to the probability of transitioning from state *i* to state *j* at time *t*). We consider a constant number of cells undergoing EMT in the system. At each point in time, a cell adopts either one of three cellular EMT phenotypes which may be epithelial, hybrid or mesenchymal. Depending on the EMT state, the cell would acquire one of *K* states of a secondary coupled process. For example, we can assume the cell cycle progression process with states *G*_1_, *G*_2_*M*, or *S* phase. We use the emission matrix *B* as follows to represent these probabilities where element *B*_*a,i*_ = Pr(*a*|*i*) represents the probability of the occurrence of state *a* of the coupled process (*a* ∈ {*C*_1_, *C*_2_, *C*_3_, …, *C*_*K*_ }) conditioned on the adoption of the *i* EMT state (*i* ∈ *E, H, M*) by the cell:

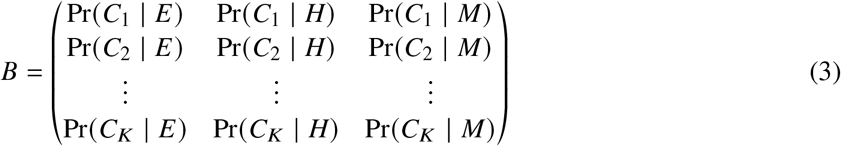

The CTMC model starting from an initial distribution of cells in each EMT state (x_0_) evolves and results in an EMT composition **x**_*t*_ . The expected observed secondary process distribution would then arise via:

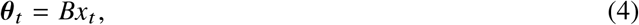

Where ***θ***_***t***_ represents the probabilities of observing each secondary process stage at time *t*, and *B* is the 3 × 3 emission matrix mapping EMT states (E, H, M) to the coupled secondary process. In Figure 1, we show a graphical representation of the framework where the EMT process comprises the latent states and the cell cycle stages are the observations.

**Figure 1.**
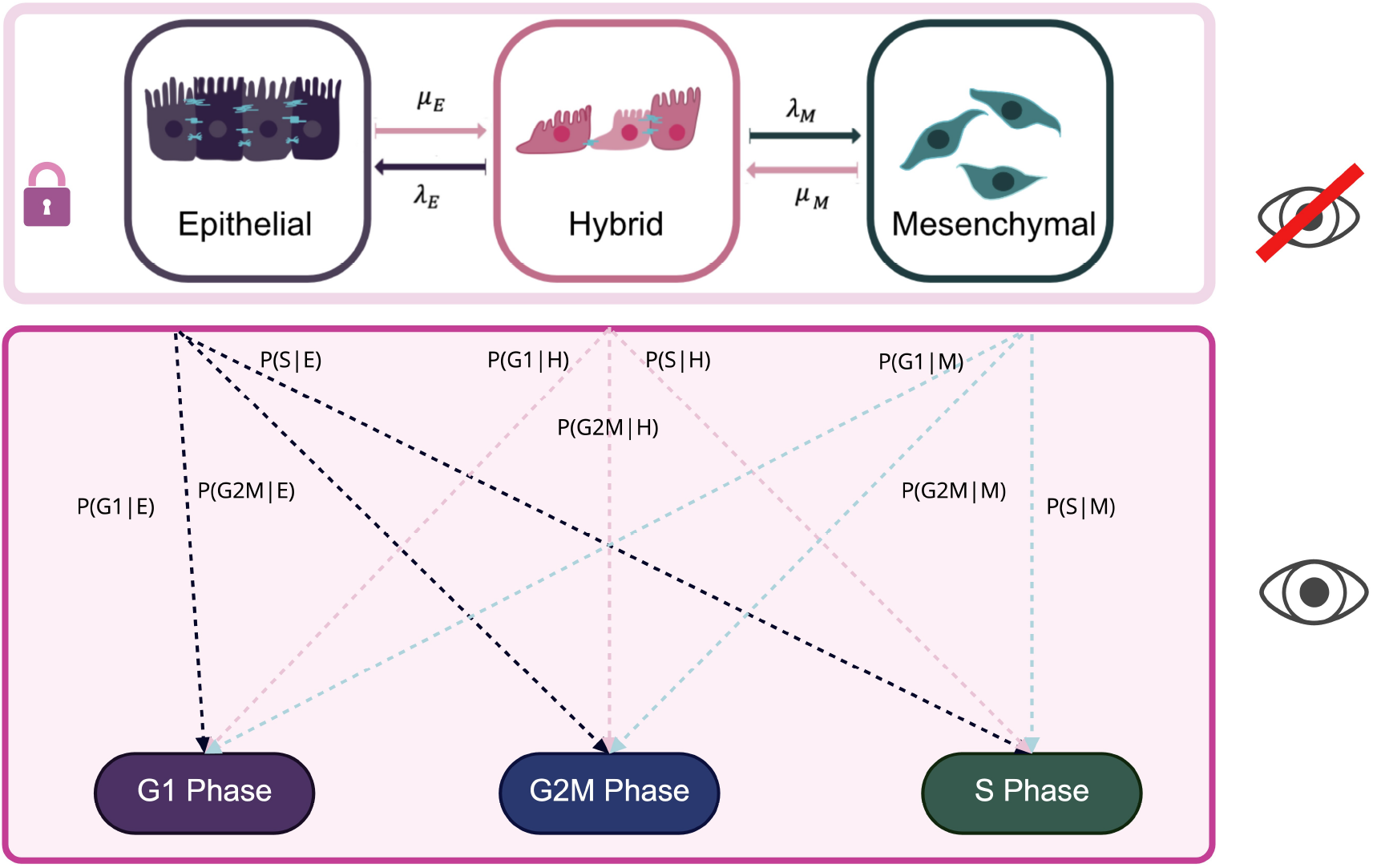
Hidden Markov Model (HMM) structure connects cell cycle observations to latent EMT states. Illustration of the HMM shows cells undergoing EMT through a CTMC with transition rates in and out of the hybrid state from or to state *i* ∈ {*E, M* } shown with *μ*_*i*_ and *λ*_*i*_ respectively (similar to the one proposed by Najafi et al (12)) and they adopt either one of three cell cycle stages: *G*_1_ phase, *G*_2_*M* phase, and *S* phase with known probabilities depending on their EMT status. We only observe the cell cycle phases and are blind to the EMT fractions. In the sections to follow we explain how we can extract the transition rates of the EMT CTMC model using cell cycle stages and the initial EMT status of the specific cell line undergoing EMT.

#### 1.1.1 Estimating the Emission Matrix

We have two sets of information:

1. **Θ**: the fraction of cells in each cell cycle stage per time point (G_1_, S, G_2*M*_ ).
2. **X**: the fraction of cells in each EMT state (E, H, M) per time point.

Our goal is to estimate the emission matrix *B* such that it satisfies the relationship:

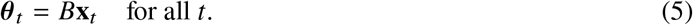

Stacking all *T* time points, we obtain the matrix form:

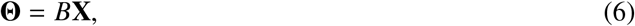

where **Θ** and **X** are 3 × *T* matrices containing the cell cycle and EMT fractions respectively across all time points.

#### 1.1.2 Least Squares Estimator

We can estimate *B* by minimizing the squared error between the observed and predicted cell cycle fractions:

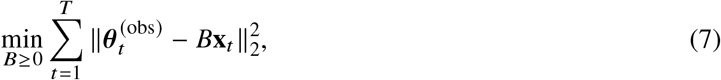

subject to the constraint that each column of *B* sums to one (since each column represents a probability distribution over all secondary process stages):

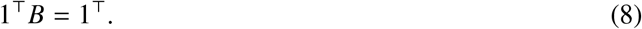

Ignoring the constraints for a moment, the closed-form least-squares solution is given by:

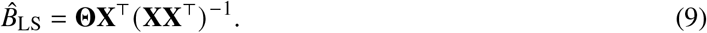

Afterward, each column of 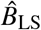 is projected onto the probability simplex to ensure non-negativity and normalization.

#### 1.1.3 Maximum Likelihood Estimator

We assume we observe only cell fractions rather than counts of cells, we interpret each observation as an empirical estimate of ***θ***_*t*_ obtained from a finite sample. Conceptually, if the total number of sampled cells at time *t* were *N*_*t*_, the observed fractions **y**_*t*_ would correspond to a multinomial sample:

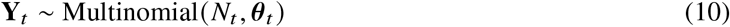

and **y**_*t*_ = **Y**_*t*_ / *N*_*t*_ would approximate ***θ***_*t*_ .

We can then estimate *B* by maximizing the log-likelihood:

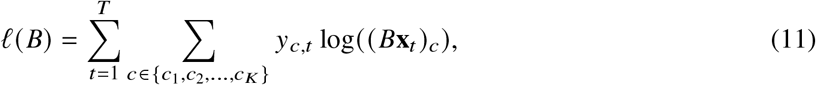

or equivalently by minimizing the negative log-likelihood:

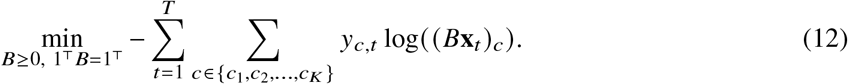

This problem can be solved using gradient-based optimization methods such as `optim()` in R, typically with the BFGS algorithm and a softmax parameterization to enforce the simplex constraints during optimization.

#### 1.1.4 Simulations

To demonstrate the ability of our framework to infer emission probabilities from two coupled processes, we ran a controlled simulation where we first fixed a ground truth emission matrix then drew EMT compositions from a Dirichlet distribution across times. Next, we computed the implied cell cycle probabilities, and we sampled multinomial counts to mimic measured data. Lastly, we re-estimated the emission matrix *B* from the aggregate data using both the closed-form least squares solution (with a post-hoc column-simplex projection) and a multinomial maximum-likelihood fit that ensures probability constraints via a column-wise softmax.

To evaluate identifiability, we swept one chosen entry of the ground truth *B* matrix (*B*_0_) across a grid of values and checked how well each estimator recovered that entry. We performed the sweep as follows: given a column *c* (one EMT state) and a row *r* (one cell cycle stage), we set the target entry *B*_*rc*_ to *v* and then renormalized the other two entries in the same column so that the column still sums up to 1. The renormalization is proportional, meaning it preserves the ratio between the untouched entries. Concretely, if the original column was (*b*_1_, *b*_2_, *b*_3_) with sum 1 and you set one probability to *v* (*b*_*r*_ = *v*), the remaining two become 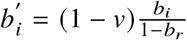, *i* ≠ *r* which guarantees nonnegativity and 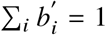 while keeping the original balance between the other categories. The function also guards edge cases such that if the original column had all its mass on the target entry (1 − *b*_*r*_ = 0), it splits the leftover 1 − *v* evenly across the other two (or equivalently uses a tiny *ϵ* to avoid division by zero). After setting that single value and renormalizing, the code regenerates data, fits *B* by LS and by MLE, and records the recovered value of that same entry. We have shown heatmaps of the results from applying LS, and MLE estimation and have compared them to the actual emission matrix in Figure 2A. As shown in the figure, both the MLE and LS approaches are able to recover the true values closely. For the simulations we used 28 timepoints, with *N* randomly drawn from 800 and 1500, and the *B*_0_ matrix below as the baseline for the parameter sweep:

**Figure 2.**
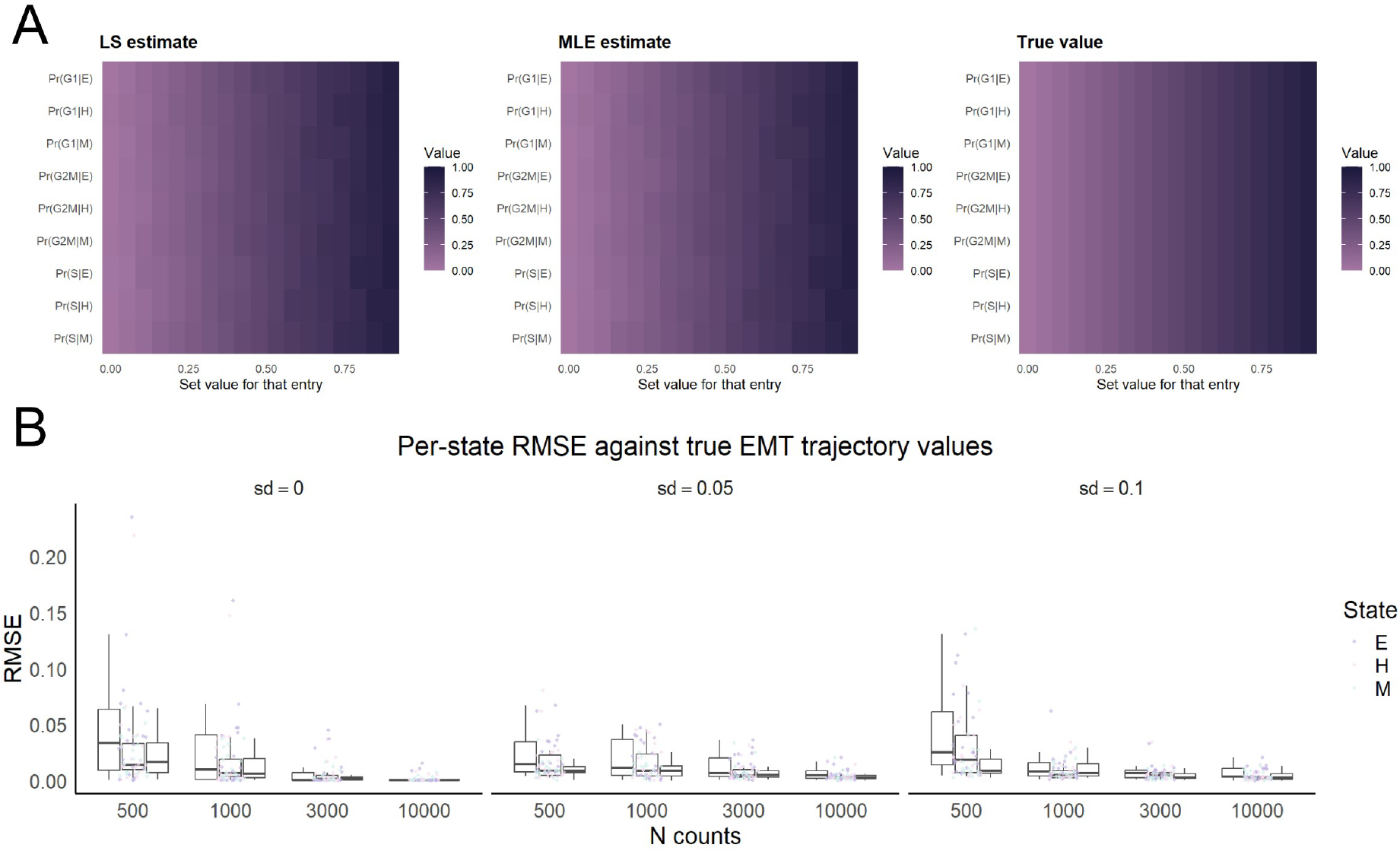
Simulations of the emission probability estimation methods and trajectory inference accuracy. A shows heatmaps of LS, MLE estimation compared to the true value of the emission matrix. B shows the RMSE calculated between the empirical EMT trajectories and simulated values obtained through HMM.

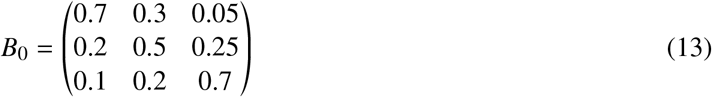

To understand how the initial sample size affects the accuracy of our estimation of the EMT trajectories given a specific *B* matrix, we performed controlled simulations. We fixed a 3-state EMT CTMC generator matrix (*Q*) and an initial distribution (*x*_0_ = [0.85, 0.1, 0.05 ]). We then propagated the hidden EMT trajectories using the transition probability matrix (*P* (*t* )) which is the matrix exponential of the generator matrix, and then generated cell cycle fractions via a known emission matrix. To mimic finite sampling, we drew multinomial counts at each time point with total sample size *N* and formed noisy fractions Θ_*obs*_ (*T* ). We repeated the analysis over different *N* values (*N* ∈ {500, 1000, 3000, 10000 }) and with a small Dirichlet noise on *B*. We have shown the distribution of the RMSE values obtained through comparing the true values from the EMT trajectories and predictions of the HMM model in Figure 2B (The script to run simulations is included within the Rmd notebooks of the Github page and can be used to test different parameter ranges). The generator matrix and emission matrices used as parameters in the simulations of Figure 2B are as follows:

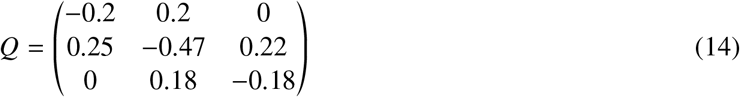

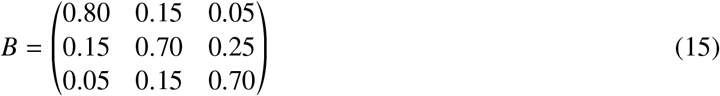

Our analysis revealed that the number of cells significantly decreases the RMSE value between predictions and true trajectory values. As *N* increases the RMSE value essentially approaches 0 when *B* is accurate. However, inaccuracy in *B* simulated via a Dirichlet noise increases the RMSE values and lowers accuracy (Figure 2B).

### 1.2 Methods

#### 1.2.1 Canonical Correlation Analysis

Canonical correlation analysis (CCA) is a statistical method that finds paired linear combinations of two sets of variables (*X* and *Y* ) which are maximally correlated with each other. CCA chooses two weight vectors *a* and *b* such that the correlation between the canonical variates *u* = *X a* and *v* = *Y b* is as large as possible, then finds subsequent pairs that are mutually uncorrelated and ordered by decreasing correlation (16). This analysis reveals shared structure across the two views which allows tasks such as exploring relationships between modalities, multi-view dimensionality reduction, and feature fusion. CCA assumes linear relationships and relies on well-conditioned covariance estimates (17). CCA is provided by default through the base `stats` package in R.

#### 1.2.2 Dynamic Time Warping Alignment

Dynamic Time Warping (DTW) alignment is an algorithm for measuring similarities between two time series data that may vary in speed or timing by nonlinearly aligning them to minimize an overall distance. It builds a cost matrix of pairwise point distances and finds the minimal-cost warping path subject to boundary, monotonicity, and step-size constraints. To perform DTW in R, we used the dtw R package (18).

#### 1.2.3 COMET

COMET which stands for context-specific optimization method of EMT trajectories was previously introduced as a method to infer EMT trajectories and inter-state transition rates from single cell RNA sequencing data. The methodology relies on the extraction of a number of highly variable EMT genes which minimize the DTW distance between the observed state-fraction trajectories and a flow cytometry ground truth data. Given the aligned trajectories, COMET fits a CTMC with generator matrix *G* to recover inter-state transition rates. COMET has several simplifying assumptions. It assumes that cells undergoing EMT eventually reach stationary distribution and that phenotypic stability factors result in symmetric transition rates into the hybrid state, both of which we relax in this study (12).

## RESULTS

### 1.3 EMT and Cell Cycle Progression Are Coupled

We started our analysis by applying our previously developed COMET R package (12, 13) to the single cell RNA sequencing dataset of Cook et al (19). We used the extracted EMT fractions and cell cycle stages as identified through Seurat (14) to proceed with the rest of our analysis. First, to understand whether the cell cycle and EMT processes are related, we performed CCA on the EMT and cell cycle stage trajectories. Our results indicated near perfect coupling of the EMT and cell cycle stage progression for the A549 and OVCA420 cell lines. As shown in Figure 3B, the coupling was to a lesser extent, but still moderate to significant for the DU145 and MCF7 cell lines. The secondary canonical variate (CV2) lines across all cell lines showed a very low to moderate *r* value which indicates that when the EMT cycle trend is removed, the remaining shared structure is very small. Overall, our analysis indicated a shared regulatory axis governing both EMT and cell cycle control which is consistent with prior studies (20). To understand whether the cell cycle progression is temporally similar across cell lines or treatment conditions, we performed pairwise DTW alignment between cell cycle stages of the different samples and have shown the results in Figure 3A. As shown in the figure, the DTW distance is lower among samples of the same cell line treated with different EMT induction factors with A549 and DU145 showing some shared cell cycle dynamics. A natural question at this stage is whether the observed coupling is inflated by shared transcriptomic signals. To test this, we compared the EMT gene set (13)with the cell cycle gene set (21) used to derive their respective trajectories and found no overlap between the two.

**Figure 3.**
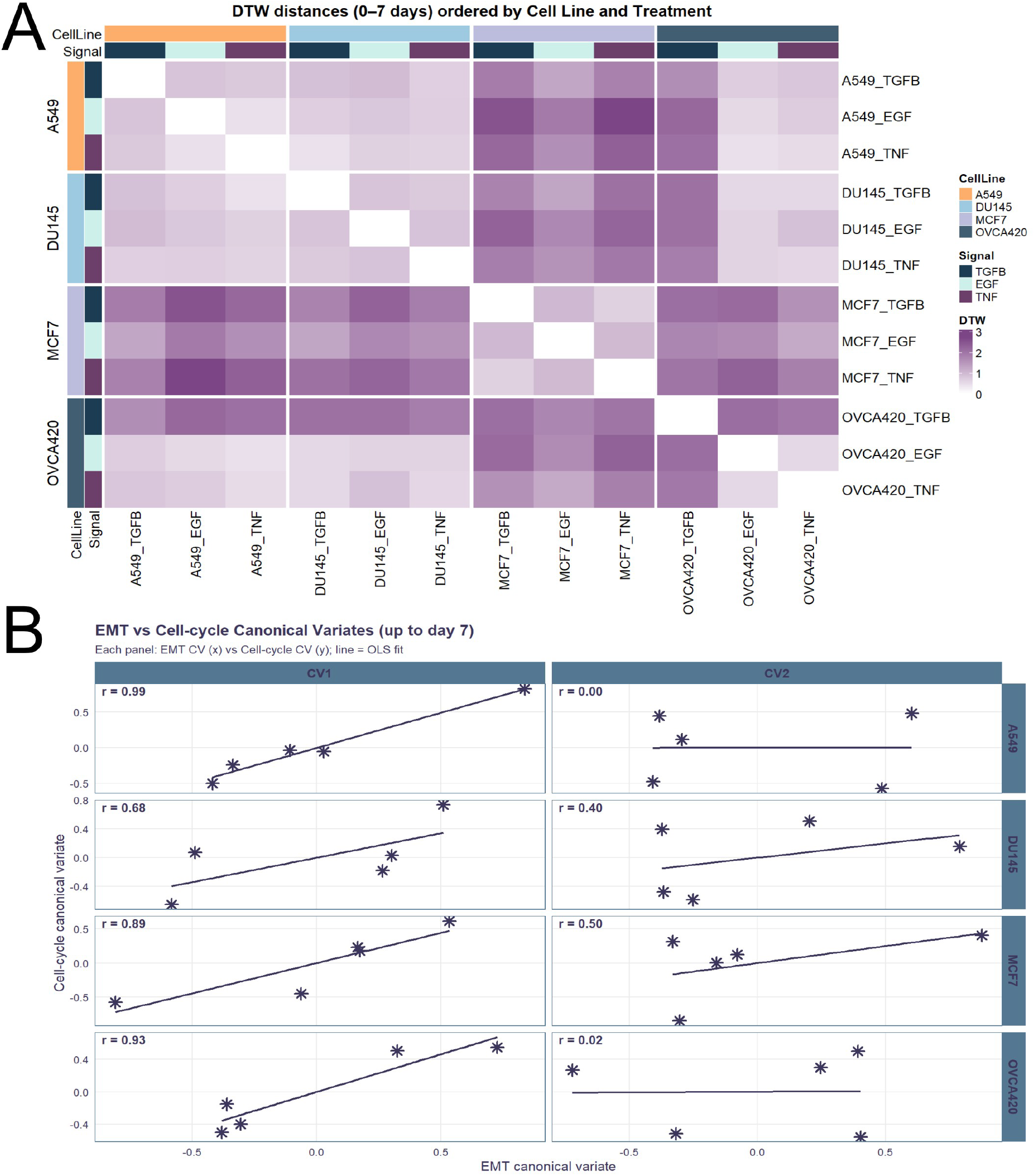
EMT and cell cycle progression are coupled. A illustrates a heatmap of pairwise DTW distances calculated on cell cycle fractions. B shows the results of CCA performed on cell cycle stage and EMT trajectories.

Overall, our findings at this stage provided a clear motivation for developing a model capable of relating the EMT dynamics and their quantitative influence on the observable cell cycle composition. In this context, the EMT states (epithelial, hybrid, and mesenchymal) can be viewed as latent populations whose transitions give rise to the observed changes in cell cycle phase distribution. The low residual correlations in the CV2 further indicated that most of the shared variance between EMT and cell cycle programs can be captured by a single dominant coupling mode, which can be effectively represented by an emission matrix linking hidden EMT states to measurable cell cycle fractions.

We note that our assumption of latent EMT states is due to the fact that while cell cycle stages can be reliably inferred from transcriptomic data using well-established marker-based classifiers, such as those implemented in `Seurat` (14) while determining EMT states remains substantially more challenging. Unlike the cell cycle process, which follows a conserved and universal transcriptional program, EMT is a context-dependent and highly plastic process with varying transcriptional signatures (19).

### 1.4 Estimating Emission Probabilities

Using the extracted EMT fractions and cell cycle stages for the TGF*β* time-course dataset from Cook et al (19), we performed MLE and LS estimation to identify the emission probability matrices for each case from day 0 up to day 7 when the EMT induction factor was withdrawn. The LS estimator minimizes the discrepancy between the observed cell cycle distributions and those predicted from the EMT fractions, assuming a linear mapping between the two processes. Formally, it solves the below equation as explained in greater detail in the Methods section:

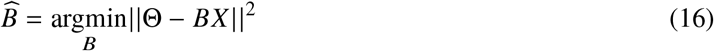

where *X* represents the EMT-state fraction matrix over time, and Θ denotes the corresponding cell cycle fraction matrix. The analytical solution to this Euclidean distance minimization is given by:

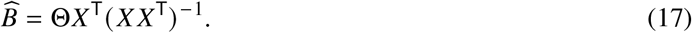

Each column of the estimated matrix 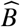 is subsequently projected onto the probability simplex (similar to the method described in Duchi et al (22)) to ensure that all entries are non-negative and that each column sums to one, thereby enforcing probabilistic consistency. This approach identifies, for each cell line, the optimal set of emission probabilities that best describe how each EMT state (epithelial, hybrid, and mesenchymal) manifests in terms of the cell cycle composition (*G*_1_, S, *G*_2_*M*) across the observed time course. The resulting matrices thus provide a quantitative, cell-line specific representation of how hidden EMT dynamics give rise to the observed proliferative behavior of the population.

In contrast, the MLE approach estimates the emission probabilities by directly maximizing the likelihood of observing the empirical cell cycle distributions given the EMT-state fractions under a multinomial probability model. Specifically, at each time point *t*, the observed cell cycle fractions 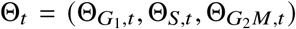 are assumed to arise from a multinomial distribution with parameters *N*_*t*_ (the number of sampled cells) and probabilities *BX*_*t*_, where *X*_*t*_ is the vector of EMT-state fractions at time *t*. The log-likelihood function for all time points is therefore given by:

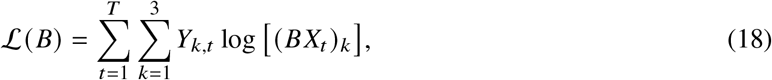

where *Y*_*k,t*_ = *N*_*t*_ Θ_*k,t*_ denotes the pseudo-counts corresponding to the observed cell cycle fractions. The MLE seeks the matrix *B* that maximizes this log-likelihood, or equivalently minimizes the negative log-likelihood:

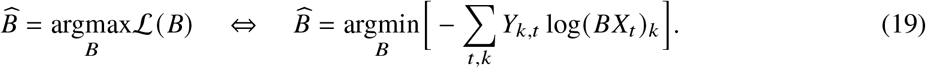

To ensure that each column of *B* remains a valid probability distribution (non-negative and summing to one), we parameterize each column as the softmax of an unconstrained three-dimensional logit vector. This reparameterization allows unconstrained numerical optimization. By iteratively adjusting these logits to maximize ℒ (*B*), the MLE procedure identifies the emission probabilities that make the observed cell cycle phase compositions most likely, given the corresponding EMT-state distributions. This probabilistic framework thus provides a statistically principled alternative to least-squares estimation, and accounts for the compositional nature of the data and the inherent constraints of multinomial sampling. To perform MLE we used the base `optim()` function from the `stats` package with the BFGS optimization method in R.

We have shown the extracted emission probability matrices found through both LS and MLE in Figure 4A and B respectively. To perform emission probability matrix extraction, we applied COMET (13) on the TGF*β* data set of Cook et al (19) with 10 runs and proceeded with the mean value for the trajectories. Across estimators, emission probabilities were very consistent with MCF7 showing the most discrepancies among its emission probabilities between the two methods. MCF7 showed a strong *G*_1_ commitment when in the epithelial state, whereas A549, DU145, and OVCA420 exhibited extremely to relatively low *G*_1_ allocation in the epithelial state. In addition to commitment to the *G*_1_ phase during the epithelial state, the MCF7 cell line also showed a high probability of acquiring the *G*_1_ cell cycle fate when in a mesenchymal state. The acquisition of the *G*_1_ phase by the MCF7 cell line is consistent with some previous studies reporting cell cycle arrest of MCF7 in *G*_1_ shortly after TGF*β* treatment (23). We previously noted that we were unsure whether this cell line actually went through EMT as a result of TGF*β* treatment or that all three EMT states were present in the original data for accurate inference of EMT trajectories (12). We note that the cell cycle fractions for this cell line as shown in our previous study are highly skewed towards the *G*_1_ phase and the *G*_1_ fraction increases over time as the cell line undergoes EMT. Overall our analysis revealed that a higher proportion of cells acquire the *G*_2_*M* and *S* cell cycle stages when in the epithelial state for all cell lines except for MCF7 which is consistent with prior studies that note active proliferation for epithelial cells compared to a lower growth rate of mesenchymal cells (7). In both our previous analysis and the present reanalysis, the EMT trajectories for OVCA420 converged toward the hybrid and mesenchymal states. The associated emission probabilities revealed elevated *G*_1_ phase occupancy for these states, consistent with prior studies reporting TGF*β*-induced G1 cell cycle arrest in ovarian carcinoma cells (24) (We note that our analysis was performed with the assumption of three stable EMT states and as explained before there is a lack of consensus on the accurate number of intermediate states in EMT (25)).

**Figure 4.**
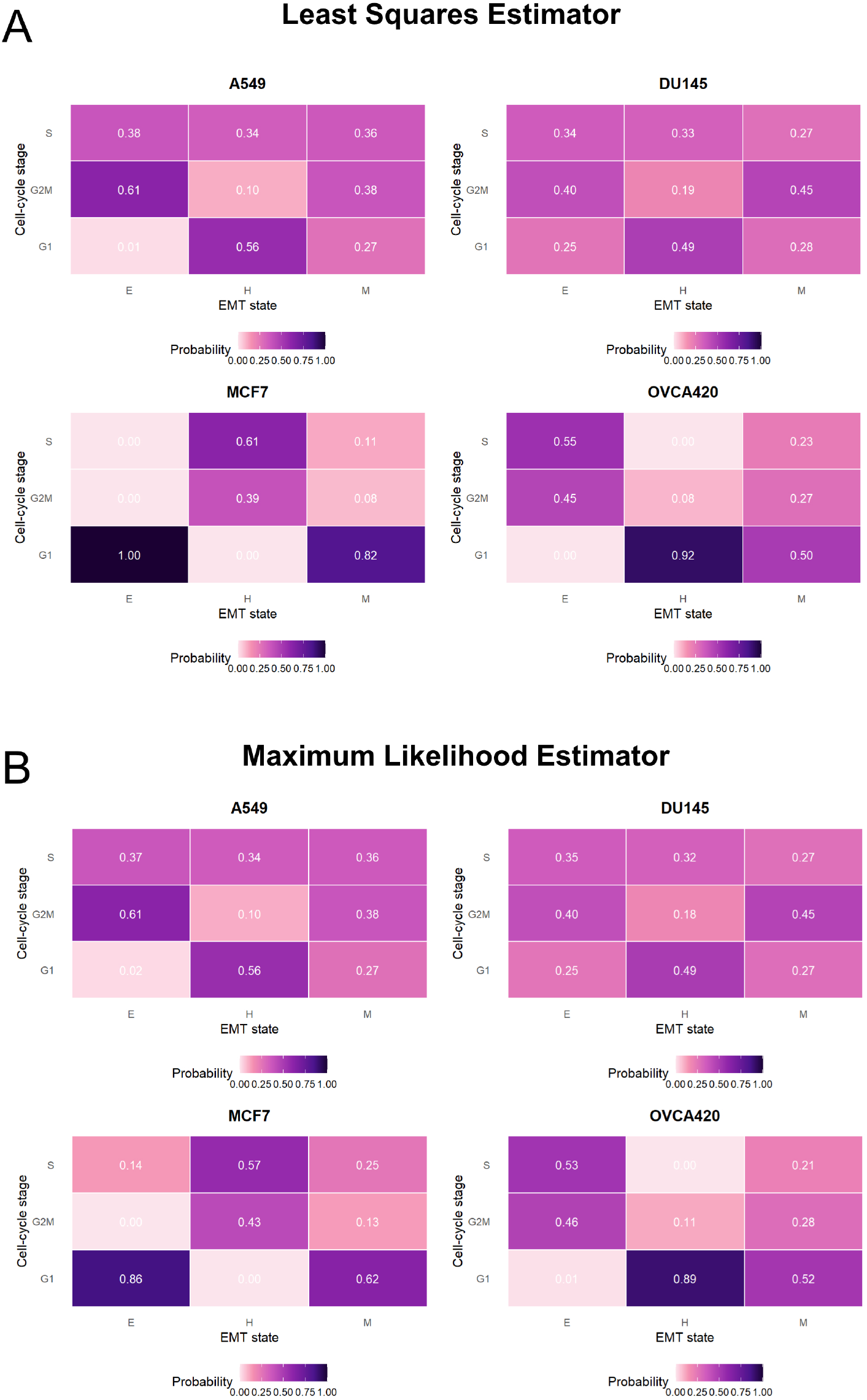
Emission probability matrices for the HMM model. A and B show the emission matrices for the HMM models obtained through Least Squares estimation and Maximum Likelihood estimation for cell lines undergoing EMT given TGF*β* treatment respectively.

### 1.5 Estimating EMT Phenotypic Distribution Based on Cell Cycle Fractions

Given our extracted emission probability matrices, an important question to address is whether we can reliably reconstruct the temporal evolution of the EMT process using only the available cell cycle stage data, a known emission probability matrix describing their coupling, and the initial EMT-state distribution of the cell line prior to treatment, which is logically assumed to remain constant. Consequently, we first proceeded by using the extracted *B* matrix to reconstruct the EMT trajectories of the TGF*β* cases of the Cook et al (19) dataset. First, we fixed the initial EMT composition *x*_0_ to the earliest pre-treatment snapshot and treated EMT progression as a three-state CTMC with a time-homogeneous generator *Q*. For each dataset, we optimized the rates of *Q* (subject to positivity and row-sum constraints) such that the model-predicted cell cycle fractions 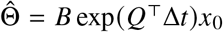 best matched the observed Θ(*t*) across time, using a multinomial (pseudo-count) likelihood. This procedure yielded a full latent EMT trajectory from cell cycle data. As shown in Figure 5 the predicted EMT trajectories shown with dashed lines match the observed EMT fractions shown with solid lines closely, except for the MCF7 cell line. With these findings, we decided to continue with testing the analysis on new data where EMT is in fact challenging to infer and EMT fractions are unavailable.

**Figure 5.**
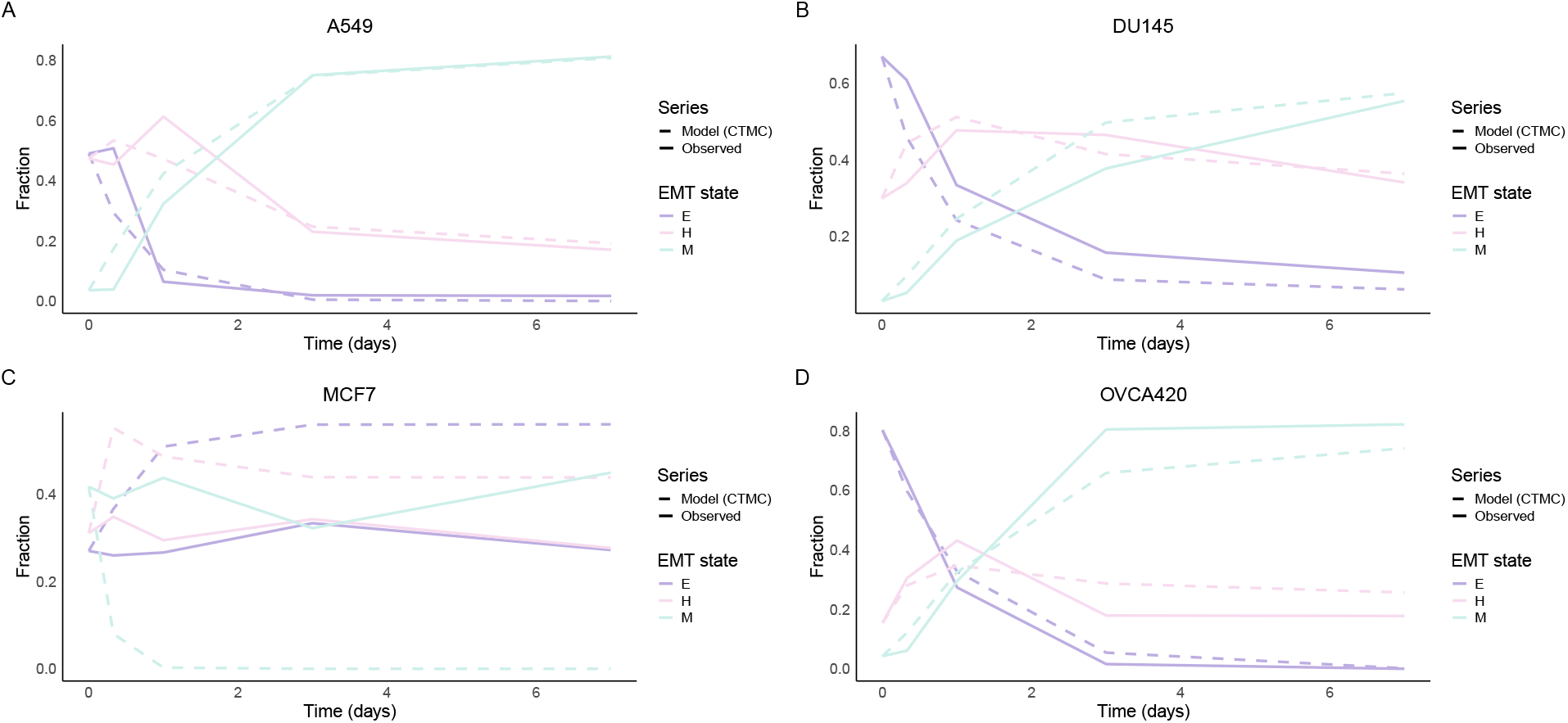
HMM can successfully reconstruct empirical EMT trajectories from cell cycle stage fractions. CTMC predicted trajectories fitted to timecourse data of cell lines undergoing EMT through treatment with TGF*β*. Dashed and solid lines correspond to model predictions and empirical observations respectively.

Because EMT and cell cycle programs are molecularly coupled (26), we expect the emission probability structure *B* to remain cell-line specific rather than treatment specific. In other words, for a given cell line, the relationship between EMT state and cell cycle phase composition should remain approximately constant even when EMT is induced by different factors, while the induction factor primarily modulates the temporal dynamics (transition rates *Q*) rather than the state-specific emissions. We note that this is a simplifying assumption we make due to lack of available sequencing data and the results shown in Figure 3A (which validate our assumption) and needs to be studied in greater detail in the future. To address this, we used a population-level HMM framework in which EMT progression is a CTMC over the three states (epithelial, hybrid, and mesenchymal), and the observed cell cycle composition at each timepoint is generated by the emission matrix *B* emanating from the latent EMT state distribution. With the initial EMT distribution *x*_0_ fixed from the baseline sample and *B* fixed from the TGF*β* data obtained previously for each cell line, we estimated the CTMC generator *Q* by maximizing the multinomial likelihood of the observed cell cycle fractions given the model predictions (Figure 6B). This yielded time-resolved EMT trajectories *X* (*t* ) inferred solely from cell cycle measurements (Figure 6A) and rate parameters that summarize the kinetics of EMT for that cell line. We have reported the corresponding four EMT interstate transition rates in Figure 7.

**Figure 6.**
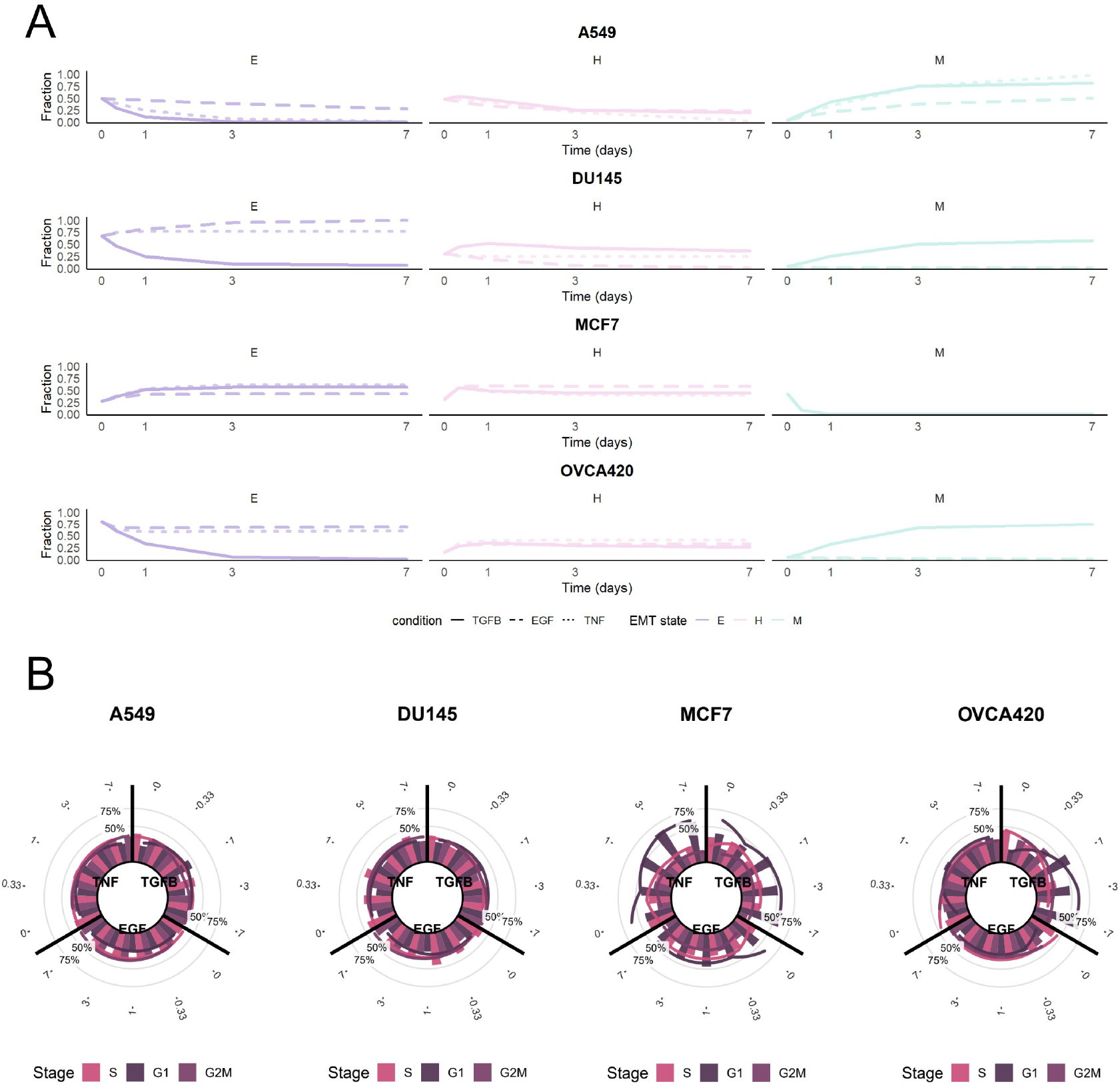
EMT trajectories of cell lines undergoing treatment with EGF and TNF*α* inferred using our HMM framework. A shows EMT trajectories for each cell line undergoing treatment with EMT induction factors with the TGF*β*, EGF, and TNF*α* induction factor treatments shown with solid, dashed, and dotted lines respectively. B shows circular plots of cell cycle fractions - percentages plotted as bars side by side with different colors as indicated in the legend - with fitted trajectories shown with relevant colors overlaid on top of the observed cell cycle fractions.

**Figure 7.**
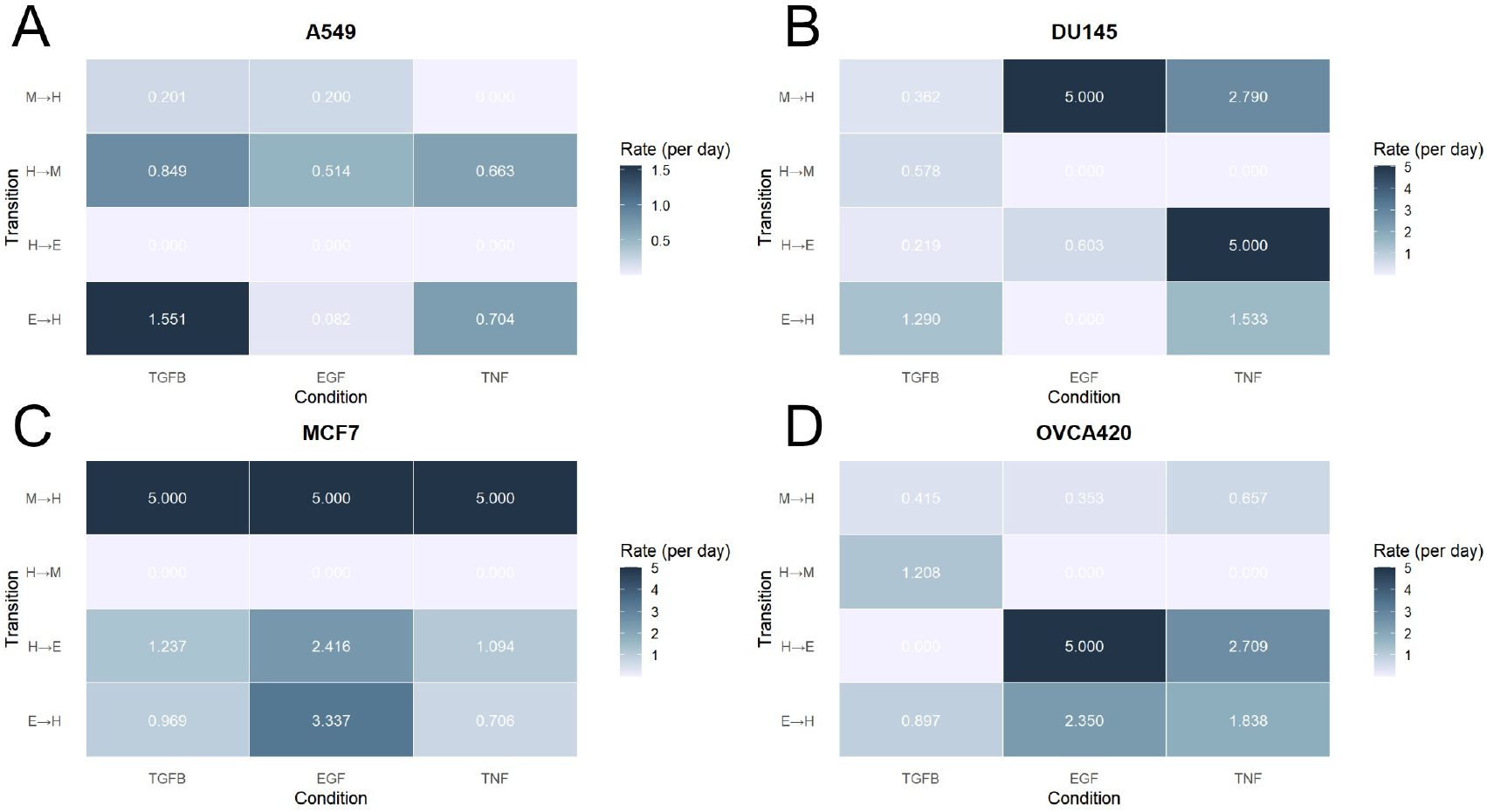
Transition rates of the CTMC model obtained through the HMM framework from cell cycle stages. A-D heatmaps show the four transition rates of the CTMC model calculated for each EMT induction factor for cell lines: A549, DU145, MCF7, and OVCA420 respectively.

As shown in Figure 6A, the A549 cell line showed significantly reduced EMT dynamics under EGF treatment. This is consistent with prior reports of EGF primarily stimulating cell migration without EMT gene expression changes as observed under TGF*β* treatment (27). On the other hand, EMT progression in the A549 cell line under TNF*α* treatment appeared to show decreased and increased transition from the epithelial to the hybrid, and hybrid to the mesenchymal state respectively. The MCF7 cell line trajectories appeared to show an increase in the hybrid state over time for all three EMT induction factors. Some studies find that MCF7 cells retain E-cadherin even after EMT induction, exhibiting only a hybrid phenotype (28). The DU145 and OVCA420 both showed significantly reduced EMT dynamics under treatment with the EGF and TNF*α* induction factors. Overall, our inferred trajectories highlight significant context-specificity. In Figure 6B we have shown the model cell cycle stage fractions overlaid on the empirical cell cycle fractions. As shown in the figure, our framework has accurately fitted cell cycle trajectories to empirical data.

As shown in Figure 7 EMT transition rates varied markedly across the four cell lines and three induction factors which highlights the context-specific nature of these state transitions (19) (we note that during fitting we constrained interstate rates to [1e-5, 5] via box-constrained optimization to avoid degenerate or implausibly fast solutions. Several estimates reached the upper bound (5/day), which indicates those parameters are boundary-constrained rather than precisely identified at our sampling resolution (0, 0.33, 1, 3, 7 days). Because the upper bound implies a median dwell time of *ln*2/5 which is around 3.3 hours and faster than our smallest interval (around 8 hours), such values should be interpreted as greater than or equal to the upper bound rather than exact rates. Likewise, values displayed as 0.000 reflect the lower bound and should be interpreted as approximately near 0. Model predictions shown in Figures 6 are generated with the same fitted generator matrix. For several transitions, the maximum-likelihood estimates reached the optimization bound which indicates that faster transition rates cannot be distinguished at our time resolution). A549 cells underwent a robust EMT under TGF*β* and displayed high rates of epithelial to hybrid, and hybrid to mesenchymal transitions. Both rates decreased under EGF and TNF*α* as expected. Compared to A549 and OVCA420, DU145 cells showed overall lower transition rates from epithelial to hybrid and hybrid to mesenchymal. This finding is consistent with prior studies that show weaker EMT induction of DU145 compared to A549 and OVCA420 (29). OVCA420 cells showed substantial EMT progression under TGF*β* treatment while undergoing little to no EMT transition under EGF or TNF*α* treatment. One surprising observation was the high transition rate from the hybrid state to the epithelial state of OVCA420 and DU145 under treatments with EGF and TNF*α* respectively. These high transition rates suggest transient EMT changes that may need further investigation. Overall, the results showed that our framework is capable of extracting interstate transition rates from cell cycle data through an HMM model. Due to lack of data, we utilized the emission matrix from the TGF*β* treated condition and shared it across other EMT inducers. While this simplification may not capture inducer-specific nuances in cell cycle coupling, it allowed for estimation of transition rates with limited measurements. Future work with inducer-specific emissions or multimodal integration is needed to further refine this approach.

### 1.6 Software Availability and Application

To enhance the accessibility of the method we proposed in this manuscript and make it available to public, we have created an R package and a web based application `hiddenCOMET` which can be used to either extract an emission probability matrix given available EMT states and cell cycle stages or reconstruct the EMT trajectories when inference is challenging given available cell cycle stages. The following code can be utilized to install the R package:

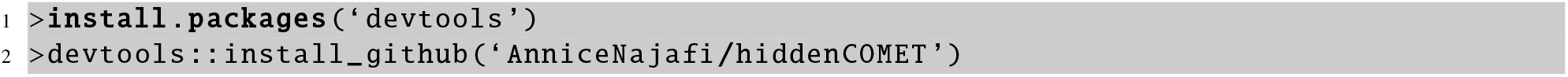

Listing 1: Installation of the R package through Github

Further instructions on how to run the code on an example are given on the Github page of the R package.

#### 1.6.1 User Interface

To access the web application, users can use the following link: https://najafiannice.shinyapps.io/hiddenComet/

The web application allows users to upload their EMT data paired with data from a secondary process to extract an emission matrix through the “Estimate B Matrix” tab on the platform. This functionality allows researchers to empirically determine the probabilistic relationships between EMT states and any number of observed cellular states, such as cell cycle phases, differentiation markers, or metabolic states.

The web application provides the ability to use two statistical methods for emission matrix estimation: least squares regression and maximum likelihood estimation. Users can upload time-course data where the first column contains timepoints (with column name ‘timepoints’) and subsequent columns represent the proportions of cells in each observed state at each timepoint, while simultaneously providing EMT trajectory data with corresponding timepoints showing the fractions of epithelial, hybrid, and mesenchymal cells. This extracted emission matrix can then be used as input for the main CTMC fitting algorithm to infer transition rates between EMT states and allow comprehensive modeling of EMT dynamics when direct estimation of the EMT distribution is challenging.

#### 1.6.2 Estimating Half Life

The app provides the functionality to calculate the half-life of each phenotype. The half-life of state *i* is calculated as follows using the waiting times obtained from the elements of the generator matrix (30):

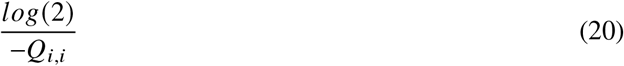

We have included the option to load the data used in this manuscript in the web-based application as demo and run the same steps outlined in this article.

## DISCUSSION

EMT remains central to metastatic progression and multiple physiological programs (26). In this study, we asked whether EMT dynamics can be inferred indirectly even when direct EMT-state calling is unreliable by exploiting the observed structure of cell cycle compositions. Building on our earlier stochastic framework (COMET) that estimates EMT trajectories from single-cell RNA sequencing (12, 13), we found that direct EMT inference can be challenging under noncanonical inducers and cell-line heterogeneity. By contrast, cell cycle stage calling is comparatively robust and prevalent (31) which prompts a complementary strategy to use cell cycle fractions as an informative readout of latent EMT states.

Our results support three main conclusions. First, EMT and cell cycle coupling is strong but cell-line specific, consistent with the context dependence reported in prior work (12). Second, an emission model learned from paired EMT/cell cycle data may potentially generalize across inducers within a cell line and enable the recovery of EMT trajectories from cell cycle fractions alone when EMT signatures are ambiguous. Third, a simple population-level HMM with a three-state CTMC provides an interpretable, quantitative link between unobserved EMT transitions and observed cell cycle fractions, and yields transition rate estimates that summarize induction-specific kinetics.

These conclusions matter because they offer a practical route when direct EMT inference fails. Our new framework allows us to salvage EMT information through a coupled process that is easier to quantify rather than discarding such datasets. Methodologically, this reframes EMT trajectory inference as a latent-state system identification problem where we can learn an emission matrix from settings where both layers are observed, then fit the CTMC generator from cell cycle fractions using the emission layer when direct EMT inference is challenging. In our analyses, This approach reproduced EMT timing and directionality in the same dataset used to estimate emissions (12).

However, our formulation is minimal and should be expanded in the future. We modeled EMT as a three-state CTMC with time-homogeneous rates. This makes parameters identifiable from modest time courses and simplifies comparisons, but it also imposes several limitations. As the field lacks consensus on the number and stability of hybrid phenotypes (2, 25), collapsing intermediate states to a single hybrid state may blur finer-grained or branching routes. Additionally time invariant generator matrix cannot capture acute to adaptation phase shifts or inducible rate changes that may happen in in-vitro EMT induction experiments (32). Lastly, our population model assumes a constant cell number, and it does not yet incorporate differential growth or death rates across EMT states. In future studies, we will add phenotypic growth rates to the generator matrix and simulate the process through a multi-type branching process. That setup will allow us to estimate the probability of extinction under different treatment conditions which would be of significant therapeutic value.

Due to the scarcity of public single cell RNA sequencing EMT datasets with longitudinal sampling, we were unable to further validate our findings. The other existing resources that we found were limited to dose-dependent responses (33) and profiles limited to a few timepoints (34) which lack the temporal structure our framework requires. Future work should validate the results of our computational framework using more time-resolved datasets across multiple cell lines and EMT induction factors.

In this study, we estimated the emission matrix *B* from the TGF*β* paired EMT/cell cycle data and, in the main analysis, held *B* fixed across inducers while fitting the EMT generator *Q*. Because *B* maps hidden EMT states to observed cell cycle fractions, allowing inducer-specific *B* is a natural extension to the current framework, but it typically requires more data. In future work we will obtain richer paired measurements and investigate this assumption further.

One additional limitation of our model is the fact that this framework is conceptualized for a system consisting of only cancer cells undergoing a biological phenomenon. It is unclear whether the direct coupling of EMT and cell cycle works in the presence of stroma and immune cells and further studies are required to understand how such cellular heterogeneity might impact the inferred transition rates or confound the emission patterns. In particular, stromal fibroblasts and infiltrating immune cells can exhibit overlapping cell cycle profiles or EMT-like gene expression, potentially introducing ambiguity in assigning observed fractions to specific cancer cell states (35, 36).

Although we utilized our model to infer EMT trajectories from cell cycle data, the framework and software provided in the R package can be applied to any dataset with a similar structure and time-resolved measurements of compositional data. This includes other differentiation processes, disease progression states, or signaling pathway activations where latent state dynamics may govern observable proportions.

## CONCLUSION

Here, we introduced hiddenCOMET, a framework for inferring EMT trajectories from single-cell sequencing data where direct inference of EMT trajectories is daunting. By leveraging the coupling between EMT and cell cycle states through a hidden Markov modeling approach, our method enables the extraction of transition dynamics from more accessible measurements such as cell cycle stages. We demonstrated that our framework recovers biologically meaningful trajectories across multiple cancer cell lines and treatment conditions, and its structure allows application to other latent processes linked to observable markers. Overall, hiddenCOMET provides an open-source statistical framework to map hidden biological programs from observable processes.

## DATA AVAILABILITY

All of data and scripts for analysis and to generate the results in the manuscript are hosted on our Github repository: github.com/AnniceNajafi/hiddenCOMET. For any questions regarding the analysis please do not hesitate to contact the corresponding author at.

## AUTHOR CONTRIBUTIONS

AN conceptualized the study, developed the methodology, performed research, created the software, analyzed results, and wrote the manuscript.

## ACKNOWLEDGMENTS

All analyses were performed in R. The figures were generated using R and assembled using Affinity Designer. Figure 1 was generated using Affinity Designer and BioRender. No funding was used to conduct this study. The software is provided free of charge and under a permissive license for public use. The authors declare no conflict of interest.

